# Reversible tuning of membrane sterol levels by cyclodextrin in a dialysis setting

**DOI:** 10.1101/2024.09.28.615506

**Authors:** Cynthia Alsayyah, Emmanuel Rodrigues, Julia Hach, Mike F. Renne, Robert Ernst

## Abstract

Large unilamellar vesicles (LUVs) are popular membrane models for studying the impact of lipids and bilayer properties on the structure and function of individual membrane proteins. The functional reconstitution of transmembrane in liposomes can be challenging, especially, if the hydrophobic thickness of the protein does not match the thickness of the surrounding lipid bilayer. During the reconstitution procedure Such hydrophobic mismatch causes low yields and protein aggregation, which are exacerbated in sterol-rich membranes featuring low membrane compressibility. Here, we explore new approaches to reversibly tune membrane sterol contents proteoliposomes after their formation. Both cholesterol delivery and extraction are mediated by methyl-β-cyclodextrin in a dialysis setting, which maintains (proteo)liposomes in a confined compartment. This makes it possible to reversibly tune the cholesterol level without losing membrane material simply by placing the dialysis cassette in a new bath containing either empty or cholesterol-loaded methyl-β-cyclodextrin. Cholesterol delivery and removal is monitored with the solvatochromic probe C-Laurdan, which reports on lipid packing. Using Förster-resonance energy transfer, we show that cholesterol delivery to proteoliposomes induces the oligomerization of a membrane property sensor, while the subsequent removal of cholesterol demonstrates the full reversibility. We propose that tuning membrane compressibility by methyl-β-cyclodextrin-meditated cholesterol delivery and removal in a dialysis setup provides a new handle to study its impact on membrane protein structure, function, and dynamics.

**Statement of significance:** Generating complex, sterol-rich, biomimetic membranes for studying the structure and function of reconstituted membrane proteins is challenging. As an important step towards asymmetric, sterol-rich, complex model membrane systems, we have established a procedure to control the membrane sterol level of liposomes and proteoliposomes using methyl-β-cyclodextrin in a dialysis setup. We demonstrate the feasibility of this approach by C-Laurdan and dehydroergosterol fluorescence spectroscopy and gain control over the membrane sterol content. We explore several parameters that affect the rate of cholesterol delivery and show that the oligomerization of a membrane property sensor, which is on the unfolded protein response sensor protein Ire1, is controlled by the sterol content of the surrounding lipid bilayer.

## Introduction

Biological membranes possess collective biophysical properties such as fluidity, thickness, and compressibility, which influence the structure, oligomeric state, and function of membrane proteins (1–4). Cholesterol is one of the most abundant lipids in mammalian cells and crucially important for modulating bilayer properties. It is asymmetrically distributed across the plasma membrane and regulates membrane permeability, membrane stiffness, and phase behavior (5–10). By increasing lipid packing, it also increases the hydrophobic thickness of the lipid bilayer and reduces membrane compressibility (11).

When the transmembrane domain of a protein does not match the hydrophobic thickness of the surrounding bilayer it is driven into oligomers to minimize the energetic strain from lipid distortion (1, 12–14). Lipid scramblases and membrane protein insertases, on the other hand, use such membrane distortions to facilitate lipid exchange between the two leaflets and/or to move hydrophilic sections of transmembrane client proteins through the hydrophobic membrane core (4, 15, 16). These examples suggest an important role of membrane compressibility in regulating membrane protein function (4). Yet, it remains challenging to define the contribution of individual membrane properties to a specific transmembrane protein function for at least three reasons. Firstly, the collective bilayer properties are physically connected and interdependent. This makes it challenging to modulate one property without perturbing others (17). Secondly, most biophysical properties of biological membranes cannot be measured directly nor deduced from their (often unknown) composition (18). Thirdly, it remains challenging to reconstitute membrane proteins in complex, asymmetric, and sterol-rich membranes for a functional characterization under defined conditions. This is particularly true for proteins such as lipid scramblases or membrane property sensors that rely on membrane distortions and hydrophobic mismatch-based mechanisms for their function. Reconstituting proteins with a substantial hydrophobic mismatch in cholesterol-rich membranes causes protein aggregation, low yields, and inhomogeneous distributions of proteins and lipids in the preparation (19–22). Hence, there is an urgent need for new experimental paradigms that would facilitate studying isolated transmembrane proteins in biomimetic, sterol-rich, defined membrane environments.

Here, we make a first step in this direction. We use methyl-β-cyclodextrin (mβCD) in a dialysis setup to modulate the cholesterol content of preformed (proteo)liposomes (23–26). MβCD is a ring-shaped, hydrophilic oligosaccharide with hydrophobic cavity (6-6.5 Å) that is sufficiently large to accommodate cholesterol. This makes mβCD a perfect, water-soluble shuttle for either delivering or extracting cholesterol and other sterols (27, 28).

Using cholesterol-loaded mβCD in a dialysis setup maintains the (proteo)liposomes confined in a separate compartment and allows for a straightforward buffer exchange and quantitative removal of cyclodextrin after sterol delivery. We follow cholesterol insertion into liposomes and proteoliposomes quantitatively using C-Laurdan spectroscopy (29) and determine the rate of cholesterol delivery and removal. We demonstrate the cholesterol-dependent, membrane-based dimerization of the membrane property sensor Ire1 that monitors membrane compressibility by a hydrophobic mismatch-based mechanism (30–32). Hence, we expand the applications of cyclodextrin in membrane research by the implementation of a dialysis setup for a direct characterization of the resulting (proteo)liposomes.

## Results

We wanted to establish an easy-to-use system for manipulating the concentration of sterols in preexisting liposomes and proteoliposomes. mβCD is an excellent tool for delivering sterols to model membranes and for removing them (33, 34). However, once mβCD and (proteo)liposomes are mixed, it is not trivial to separate them. Normally, this separation involves size exclusion chromatography or harvesting the (proteo)liposomes by ultracentrifugation, which is both time-consuming and inefficient (35). We wanted to know if mβCD in a dialysis setup can support lipid exchange whilst retaining (proteo)liposomes in a separate compartment. This would make subsequent preparative steps obsolete and minimize loss of material.

Both empty and cholesterol-loaded mβCD (Fig. 1A) can cross a dialysis membrane with a molecular weight cutoff of 100 kDa (pore size ≈10.5 nm) (Fig. 1B). By dialyzing (proteo)liposomes contained in the dialysis cassette with either cholesterol-loaded or empty mβCD, we should be able to either deliver cholesterol or to remove it (Fig. 1B). Because the (proteo)liposomes are trapped within the cassette, it should be possible to repeat this procedure several times. Hence, this experimental setup provides a means to reversibly manipulate the cholesterol concentration in pre-formed (proteo)liposomes to modulate lipid packing and membrane compressibility.

**Figure 1:**
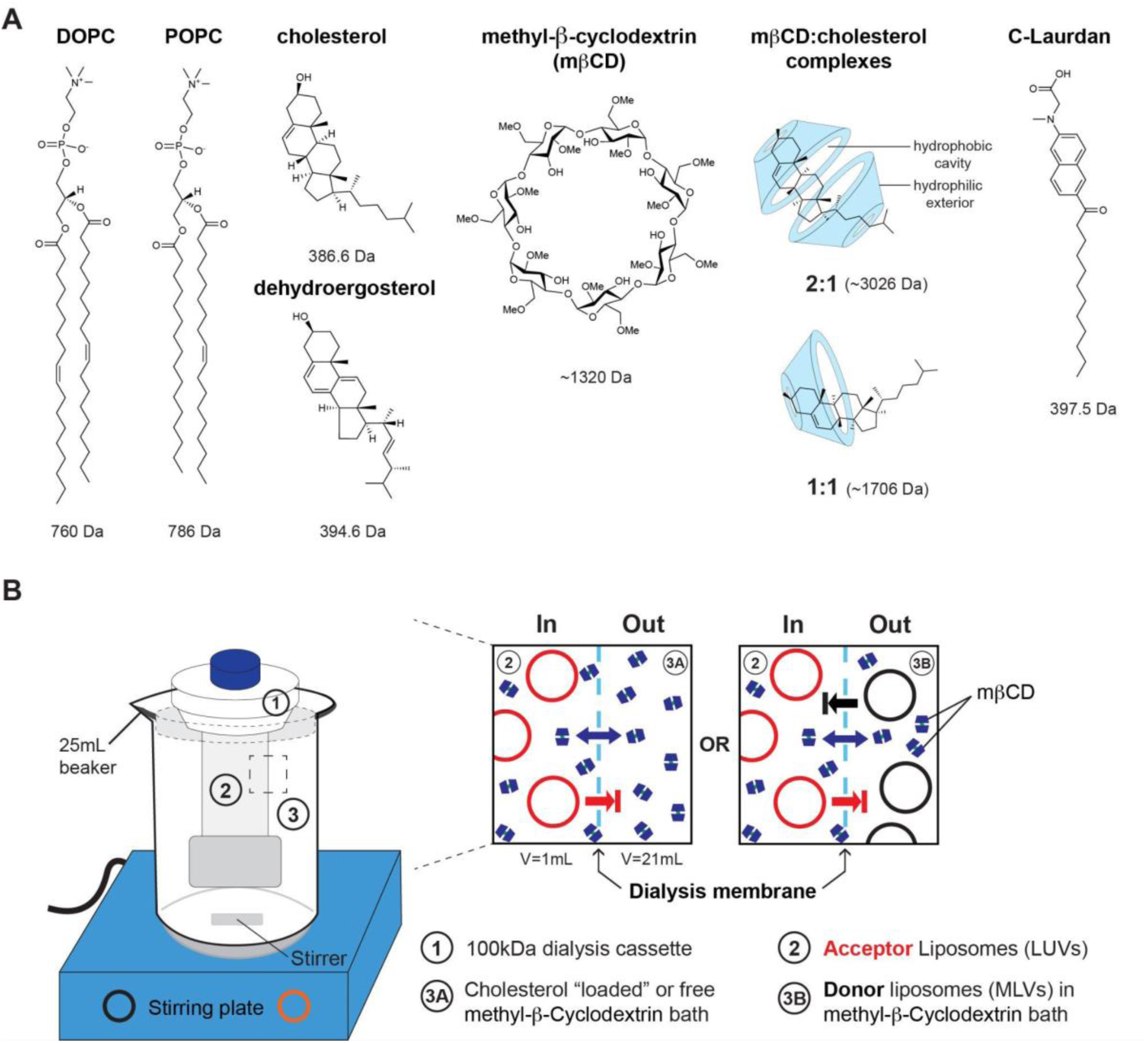
Manipulating cholesterol levels in preformed membranes. **(A)** Chemical structures of molecules used in this study. POPC: 1-palmitoyl-2-oleoyl-*sn*-glycero-3-phosphocholine; DOPC: 1,2-dioleoyl-*sn*-glycero-3-phosphocholine. mβCD is a cyclic oligosaccharide that can bind cholesterol and other sterols such as dehydroergosterol in different stoichiometries. C-Laurdan is a solvatochromic probe that reports on the degree of water penetration into a lipid bilayer. **(B)** A dialysis cassette with a molecular weight cutoff of 100 kDa (1) is filled with a suspension of (proteo-)liposomes (2). The (proteo)liposomes are trapped and separated from the outer bath containing either cholesterol-loaded or empty mβCD. Alternative setups with donor liposomes present in the outer bath are conceivable. MβCD can pass the dialysis membrane to either deliver or pick up cholesterol.

### Kinetics of cholesterol delivery through a dialysis membrane

First, we were interested in the delivery of cholesterol to LUVs composed of 1-palmitoyl-2-oleoyl-*sn*-glycero-3-phosphocholine (POPC), an abundant lipid in bio-membranes featuring 50% unsaturated monounsaturated lipid acyl chains (18). Cholesterol was delivered to the LUVs in a dialysis cassette by placing it in an outer bath (V = 21 ml) containing 2.4 mM cholesterol-loaded mβCD. Cholesterol incorporation was probed, by removing 60 µl fractions of the liposome suspension at various times points and subjecting them to C-Laurdan spectroscopy (29, 36) (Fig. 2A). C-Laurdan is a solvatochromic probe that reports on the degree of water penetration into the lipid bilayer, which is directly related to the inter-lipid spacing (29, 36, 37). The generalized polarization (GP) of C-Laurdan is a ratiometric value derived from the fluorescence emission spectrum. It can assume values from +1 (being most ordered) to −1 (being least ordered), but its absolute value depends on the instrumentation and many other factors (36). Previously, giant unilamellar vesicles (GUVs) formed from POPC featured a C-Laurdan GP of −0.29, while the inclusion of 40 mol% cholesterol resulted in a GP of 0.27 indicative for tighter lipid packing (36). In our setup, we observed a dramatic change of the C-Laurdan fluorescence emission spectrum within a few hours of cholesterol delivery (Fig. 2A). Hence, cholesterol-loaded mβCD can readily pass the dialysis membrane and unload its cargo into the membrane of acceptor liposomes. Next, we tested if cholesterol can be removed from these liposomes. Upon placing the dialysis cassette with cholesterol-containing, POPC-based liposomes in a new bath containing 4.8 mM empty mβCD and a two-fold excess of POPC-based, multilamellar liposomes as a ‘sink’ for cholesterol, we observed changes in the fluorescence emission spectrum consistent with near-complete cholesterol removal (Fig. 2B). With respect to the C-Laurdan GP values, we observed an increase from −0.15 to 0.24 upon cholesterol delivery (Fig. 2C) and a decrease back to −0.16 upon cholesterol removal (Fig. 2C). While the observed changes of the C-Laurdan spectrum suggested an efficient modulation of cholesterol in the liposome membrane, we wanted to determine its precise concentration. To achieve this, we performed a calibration experiment (Fig. S1A). We generated a series of POPC-based LUVs containing different cholesterol concentrations, recorded C-Laurdan fluorescence emission spectra, and plotted the experimentally determined GP values against the known cholesterol concentration under identical conditions (Fig. S1A). The observed dependency of the GP values from the cholesterol concentration could be fitted reasonably well with a polynomial function, thereby providing a means to deduce the molar concentration of cholesterol based on a C-Laurdan fluorescence emission spectrum (Equation 2 – Fig. S1A). Making use of this calibration, we found that the cholesterol concentration in POPC-based liposomes reaches 31.5 mol% within five hours of delivery with an approximated initial rate of 0.22±0.01 mol% min^−1^ using the derivative of a mono-exponential association function. Notably, almost all cholesterol can be removed again when dialyzing the cholesterol-loaded liposomes with empty mβCD (Fig. 2D).

**Figure 2:**
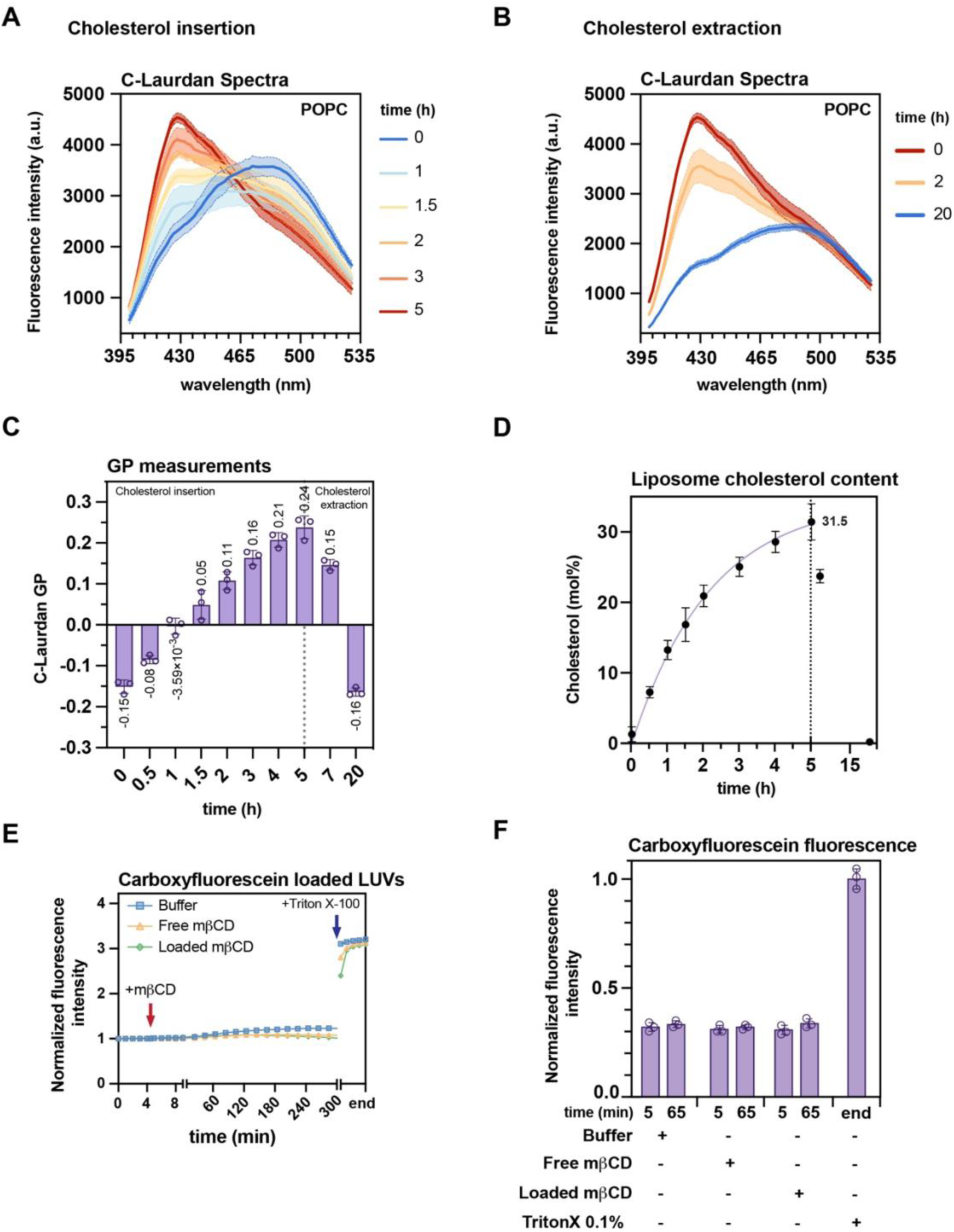
POPC-based large unilamellar vesicles remain intact during cholesterol insertion and removal. **(A)** C-Laurdan fluorescence emission spectra (Ex: 375±5 nm; Em slit width: 5 nm) of POPC-based liposomes retrieved from the dialysis cassette after indicated times of cholesterol delivery using mβCD. Data are from three independent experiments (n = 3;mean±SD). **(B)** C-Laurdan fluorescence emission spectra upon cholesterol removal using empty mβCD in the outer bath of the dialysis setup. Data are from three independent experiments (n = 3;mean±SD). **(C)** C-Laurdan GP of pure POPC LUVs during a cholesterol delivery and removal experiment over 20 h at room temperature. The dotted vertical line (at t = 5h) indicates the exchange of the outer batch and a switch from cholesterol insertion to cholesterol extraction. Data are from three independent experiments (n = 3; mean±SD). **(D)** Cholesterol concentration (in mol%) derived from a standard curve. The dotted vertical line (at t = 5h) indicates the exchange of the outer batch and a switch from cholesterol insertion to cholesterol extraction. The purple solid line fitted to the data was obtained using a one phase association model (Prism 10). Shown are data from three independent experiments (n = 3; mean±SD). **(E)** Representative carboxyfluorescein (CF) fluorescence emission (Ex: 492±10 nm; Em: 517±10 nm) in CF-loaded, POPC-based liposomes (20 µM lipids) over time. At t = 5 min (red arrow), the liposomes were either treated with 220 µM cholesterol-loaded mβCD (green), 440 µM empty mβCD (yellow) or mock untreated (blue). Addition of the detergent Triton X-100 (final concentration 0.1% (w/v)) (blue arrow) solubilizes the membrane and triggers the release of CF. **(F)** The CF emission was normalized to the maximally recorded emission after the addition of Triton X-100. Represented are two timepoints: t = 5 min (time of mβCD/buffer addition) and t = 65 min (after 60 min of mβCD/mock treatment). Data are derived from three independent experiments (n = 3; mean±SD).

The same set of experiments was performed with liposomes containing an equimolar mix of POPC and DOPC featuring 75% mono-unsaturated lipid acyl chains (Fig. S2A-E). Consistent with the lower degree lipid packing in unsaturated membranes, we also observed lower C-Laurdan GP values for DOPC:POPC-based liposomes (Fig. S2C-E) than for POPC-based liposomes (Fig. 2D,E). During dialysis with cyclodextrin, we again observed efficient delivery and removal of cholesterol (Fig. S2A-C) and found that concentrations of ≈30 mol% are reached within five hours of dialysis (Fig. S2D,E). The initial rate of cholesterol delivery k = 0.16±0.01 mol% min^−1^ was ≈1.5-fold lower (Fig. S2E) compared to the more tightly packed POPC liposomes (Fig. 2D).

After prolonged dialysis (>24 h) the system approached an equilibrium with barely any changes of the C-Laurdan GP (Fig. S2F). Notably, the final concentration of cholesterol was higher in the more loosely packed membrane containing both DOPC and POPC (52.24 mol%) than in the more tightly packed POPC-based membrane (42.65 mol%) (Fig. S2F). Our observations suggest that the loosely packed membrane has a ≈10% higher capacity for cholesterol although cholesterol has a higher affinity to POPC (38–44).

### Liposomes remain intact during cholesterol exchange

When used in excess, mβCD can dissolve liposomes (24, 45, 46). Even though the lipid:mβCD ratio of 1:11 in our setup is much lower than the 1:100 ratio required for solubilization (46, 47), we wanted to verify that our liposomes remain intact during cholesterol delivery (Fig. 2E,F). To this end, we generated POPC-based liposomes loaded with 75 mM carboxyfluorescein (CF). CF is a water-soluble, charged fluorophore that is only poorly fluorescent at high concentrations due to excimer formation and self-quenching (48). Hence, for as long as CF is contained in the liposomes, we expect the sample to exhibit only a low level of fluorescence emission. Membrane rupture or solubilization, however, should lead to unquenching of CF marked by an increase in fluorescence intensity. We recorded the CF emission during the incubation of the respective liposomes with either cholesterol-loaded or empty mβCD in real time (Fig. 2E). As a control, we also incubated the CF-loaded liposomes in a buffer lacking mβCD (Fig. 2E). During five hours of incubation, we observed only minor changes of the fluorescence emission (Fig. 2E,F) compared to the 3-fold increase observed upon solubilizing the liposomes with the detergent Triton X-100 (final concentration ≈1.6 mM) (Fig. 2F,G). We conclude that the liposomes remain intact during the incubation both with cholesterol-loaded mβCD or empty mβCD. This interpretation was corroborated by dynamic light scattering (DLS) experiments, which demonstrate that the average diameter of the liposomes (197±0.7 nm) is not affected by cholesterol delivery or by cholesterol removal (Fig. S1B-D). Furthermore, DLS experiments confirmed the efficacy of Triton X-100 in solubilizing liposomes: For detergent-treated liposomes we detected only objects of ≈13 nm, which likely represent mixed micelles of Triton X-100 and membrane lipids (Fig. S1C). Together, our experiments suggest that preformed liposomes remain intact when their cholesterol level is remodeled using mβCD.

### Modulating the cyclodextrin-dependent cholesterol exchange kinetics

Next, we explored how the temperature and the molecular weight cutoff of the dialysis membrane affects the rate of sterol transfer. Initially, we performed the delivery of cholesterol into pure POPC acceptors at three different temperatures (4°C, 22-24°C, and 55°C) and across a dialysis membrane with a 100 kDa molecular weight cutoff (Fig. 3A). For each temperature, we followed the C-Laurdan GPs during sterol delivery over time. Expectedly, the GP increased gradually for all temperatures but more dynamically at higher temperatures (Fig. 3A). Using an established calibration curve (Fig. S1A), we determined the cholesterol concentration in the acceptor liposomes for a more quantitative insight. Fitting these data to a one phase exponential association model revealed the impact of temperatures on the rate of cholesterol delivery (Fig. 3B): At 55°C we obtained a 4-fold faster initial rate of cholesterol incorporation (k_55°C_ = 0.33±0.04 mol% min^−1^) than at 4°C (k_4°C_ = 0.08±0.07 mol% min^−1^). Likewise, the molecular weight cutoff may affect the rate of cholesterol delivery to acceptor liposomes (50 mol% POPC and 50 mol% DOPC). We tested three different dialysis membranes with molecular weight cutoffs of 3.5 kDa, 20 kDa, and 100 kDa (Fig. 3C-D). Compared to the rapid delivery over a dialysis membrane with a 100 kDa molecular weight cutoff (pore size of 6-10 nm), the delivery was dramatically slower across membranes with 20 kDa and 3.5 kDa molecular weight cutoffs (Fig. 3D). The most obvious explanation is the reduced pore size, which is only 3-5 nm and 1-2 nm for the membrane with molecular weight cutoffs of 20 kDa and 3.5 kDa, respectively. Thus, the passage of mβCD across the dialysis membrane can become rate-limiting for cholesterol delivery. This is important to consider when choosing the dialysis membrane for a particular experiment. For example, a membrane with lower pore sizes could be used to ‘ramp up’ the cholesterol concentration very slowly in the (proteo)liposomes of interest.

**Figure 3:**
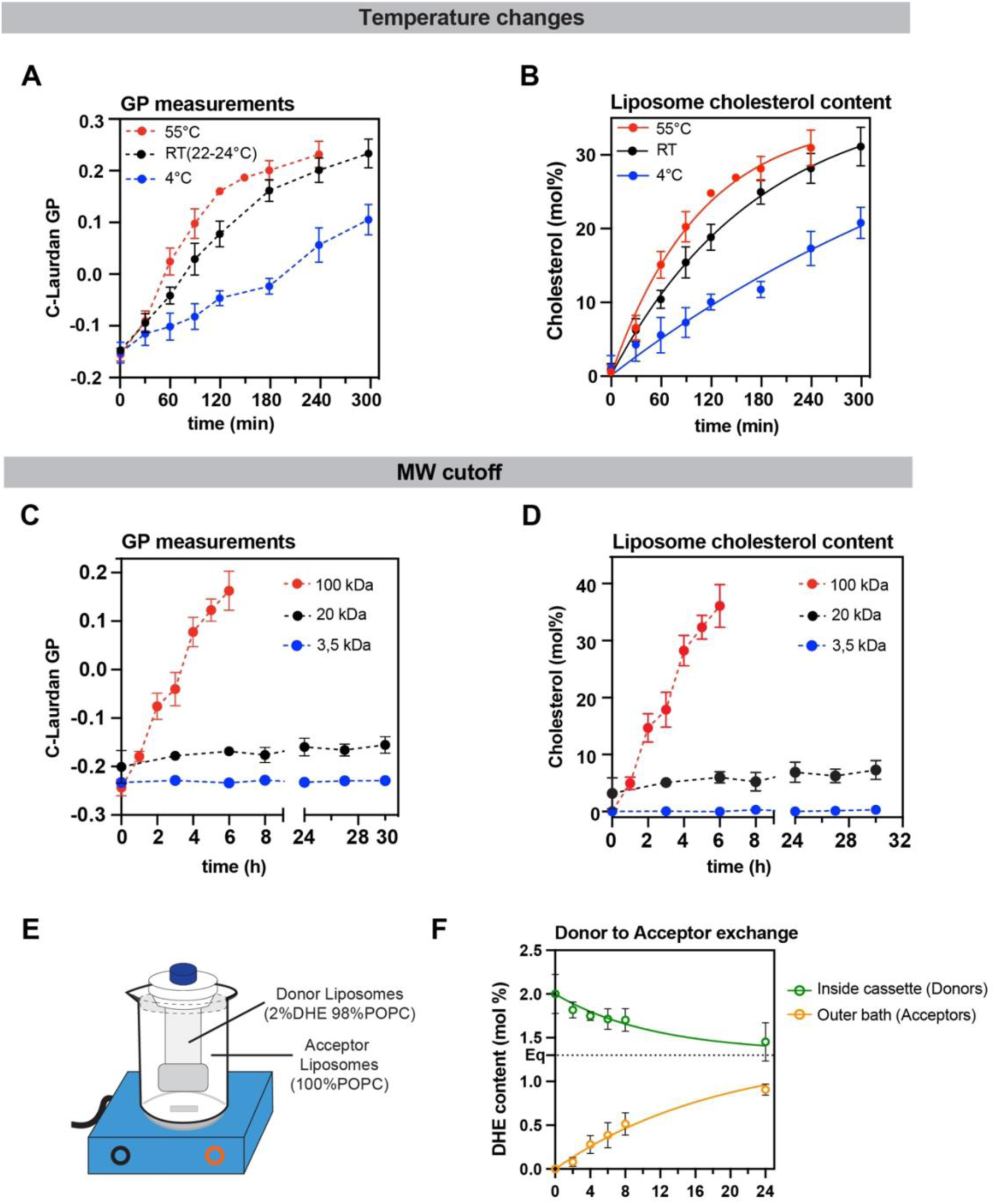
Cholesterol insertion into liposomes is modulated by temperature and the molecular weight cutoff of the dialysis membrane. **(A)** C-Laurdan GP values of POPC-based LUVs during five hours of cholesterol delivery at 4°C (blue), RT (23±1°C) (black) and 55°C (red). Data are from three independent experiments (n = 3; mean±SD). **(B)** The molar fraction of cholesterol is derived from the GP values in (A) and plotted with the same color code as in (A). The data were fitted using a one phase exponential association model. The data are from three independent experiments (n = 3; mean±SD). **(C)** C-Laurdan GP values of LUVs at various times during cholesterol delivery initially composed of 50 mol% POPC and 50 mol% DOPC. The experiment was performed with dialysis membranes differing in their molecular weight cutoffs: 3,5-5 kDa (blue); 20 kDa (in black) and 100 kDa (red). The data are derived from three independent experiments (n = 3; mean±SD). **(D)** The molar fraction of cholesterol is derived from the GP values in (C) and plotted with the same color code as in (C). **(E)** Sonicated liposomes composed of 98 mol% POPC and 2 mol% (Donors) were dialyzed in the presence of empty mβCD against a two-fold excess of POPC-based liposomes using a dialysis cassette with a 100 kDa molecular weight cutoff. **(F)** The transfer of DHE from donor liposomes (200 µmol) to acceptor liposomes (400 µM) at 23±1°C was followed using the DHE fluorescence emission (Ex: 324±4 nm; Em: 394±4 nm) yielding an estimate of the molar DHE concentration in the donor and acceptor liposomes (green and yellow, respectively). For the donor liposomes, the data were fitted using a one phase exponential decay (constraint at Y0 = 2.0 mol% DHE and plateau of 1.33 mol% DHE) with an R^2^ = 0.93. For the acceptor liposomes, the data were fitted with a one phase association model (constraints for Y0 = 0 mol% DHE and the plateau of 1.33 mol% DHE) with an R^2^ = 0.98. The data are derived from three independent experiments (n = 3; mean±SD).

### Sterol exchange between donor and acceptor liposomes

We wanted to know if the dialysis setup is suitable to transfer sterols between liposomes. To this end, we followed the concentration of dehydroergosterol (DHE) in donor and acceptor liposomes, which were separated by a dialysis membrane (Fig. 1A). DHE is a fluorescent analogue of ergosterol with three conjugated double bonds (Fig. 1A; Fig. S3A). Over a broad range of concentrations, the fluorescence emission of DHE is proportional to its molar concentration in liposomes (Equation 4 - Fig. S3A). This allowed us to directly quantify DHE in both compartments (Fig. 3E) thereby monitoring both cholesterol extraction and delivery (Fig. 3F). In the dialysis cassette we used sonicated POPC-based liposomes with 2% DHE (donor liposomes). In the outer bath, we used a 2-fold excess of liposomes composed only of POPC (acceptor liposomes), which are initially non-fluorescent (Fig. 3E). Over time, we observed a decrease in the fluorescence in the compartment with the donor liposomes and an increase in the compartment with acceptor liposomes both approaching the theoretical equilibrium concentration of 1.33 mol% (Fig. 3F). However, our approach cannot distinguish between DHE bound to mβCD and DHE in the membrane. Assuming that the majority of DHE is in the membrane under these conditions (40), we fitted the experimental data using single exponential equations for DHE removal and delivery with a forced plateau at 1.33 mol%. The initial rate of DHE removal from the donor liposomes k_don_ was −0.06±0.01 mol% min^−1^ and indistinguishable from the initial rate of delivery into the membrane of acceptor liposomes k_acc_ = 0.07 mol% min^−1^. This suggests that either DHE removal or the transfer of cholesterol-loaded mβCD through the dialysis membrane is rate-limiting. Furthermore, the dialysis setup helps monitoring changes of the sterol level in two communicating populations of liposomes.

### Studying the impact of cholesterol on the oligomerization of Ire1

Transmembrane protein reconstitution in sterol-rich membranes is challenging especially when there is a significant hydrophobic mismatch between the protein and the lipid bilayer. Energetic penalties associated with hydrophobic mismatch are higher in sterol-rich membranes (13), thereby lowering the efficiency of transmembrane protein insertion. We wanted to test if our setup can provide a means to modulate sterol content after the formation of proteoliposomes.

We decided to study a model transmembrane protein based on the membrane property sensor Ire1 from *Saccharomyces cerevisiae*. Ire1 uses a hydrophobic mismatch-based mechanism to sense aberrant stiffening and thickening of the endoplasmic reticulum membrane (ER) (4, 30, 31). Increased membrane thickness and reduced ER membrane compressibility in the cells drives Ire1 into dimers and higher oligomers, which ultimately, triggers the unfolded protein response controlling hundreds of target genes (30, 32, 49, 50). Previously, the impact of the membrane environment on Ire1 dimerization was established by continuous wave electron paramagnetic spectroscopy (cwEPR) using a spin-labeled minimal sensor protein derived from Ire1 (30). Here, we used Förster resonance energy transfer (FRET) to assess the oligomeric state of a similar Ire1-based sensor construct as an alternative readout. This Ire1-based sensor construct consists of an N-terminal Maltose binding protein, a flexible linker with a tobacco etch virus (TEV) protease recognition site, and the residues P501 to K570 from Ire1, covering its entire transmembrane region with a functionally relevant amphipathic helix and the short transmembrane helix (30, 31). A single cysteine was introduced at the C-terminal end of the construct to facilitate fluorescent labeling by maleimide-based chemistry. To prevent complications from undesired covalent crosslinking of Ire1, the endogenous cysteine 552 in the transmembrane helix was replaced by serine. Previously, it was shown that this mutation does not affect the function of Ire1 (30, 31).

We co-reconstituted a ATTO514- and ATTO594-labeled constructs in liposomes composed of 50 mol% POPC and 50 mol% DOPC at a molar protein-to-lipid ratio of 1:16000, which should minimize proximity-FRET from random encounters of labeled Ire1 molecules in the liposome membrane. The acyl chain composition was chosen to reflect the acyl chain composition of the yeast ER in both length and lipid saturation (50, 51). Under these conditions, we expect no membrane-based dimerization of Ire1 and therefore no energy transfer between the two fluorophores even though they can form a FRET pair. Indeed, upon excitation of the ATTO514-labeled donor at 514 nm, we detected barely any energy transfer that would give rise to an emission at 620 nm due to FRET (Fig. S4A).

When we used our dialysis setup to deliver cholesterol to the FRET pair-containing proteoliposomes using mβCD, we observed over time increasing FRET signals (Fig. 4B,C) suggesting an increased proximity of the donor and the ATTO594-labeled acceptor constructs likely caused by the dimerization of Ire1. No such changes in the fluorescence spectrum were observed in proteoliposomes containing the ATTO514-labeled donor alone (Fig. S4C). The increased FRET signal upon cholesterol delivery to FRET pair-containing liposomes was abolished, when the proteoliposomes were solubilized with detergents (Fig. S4B-D). We conclude that successful cholesterol delivery triggers a change in membrane properties that induces dimerization/oligomerization of Ire1-derived sensor constructs.

**Figure 4:**
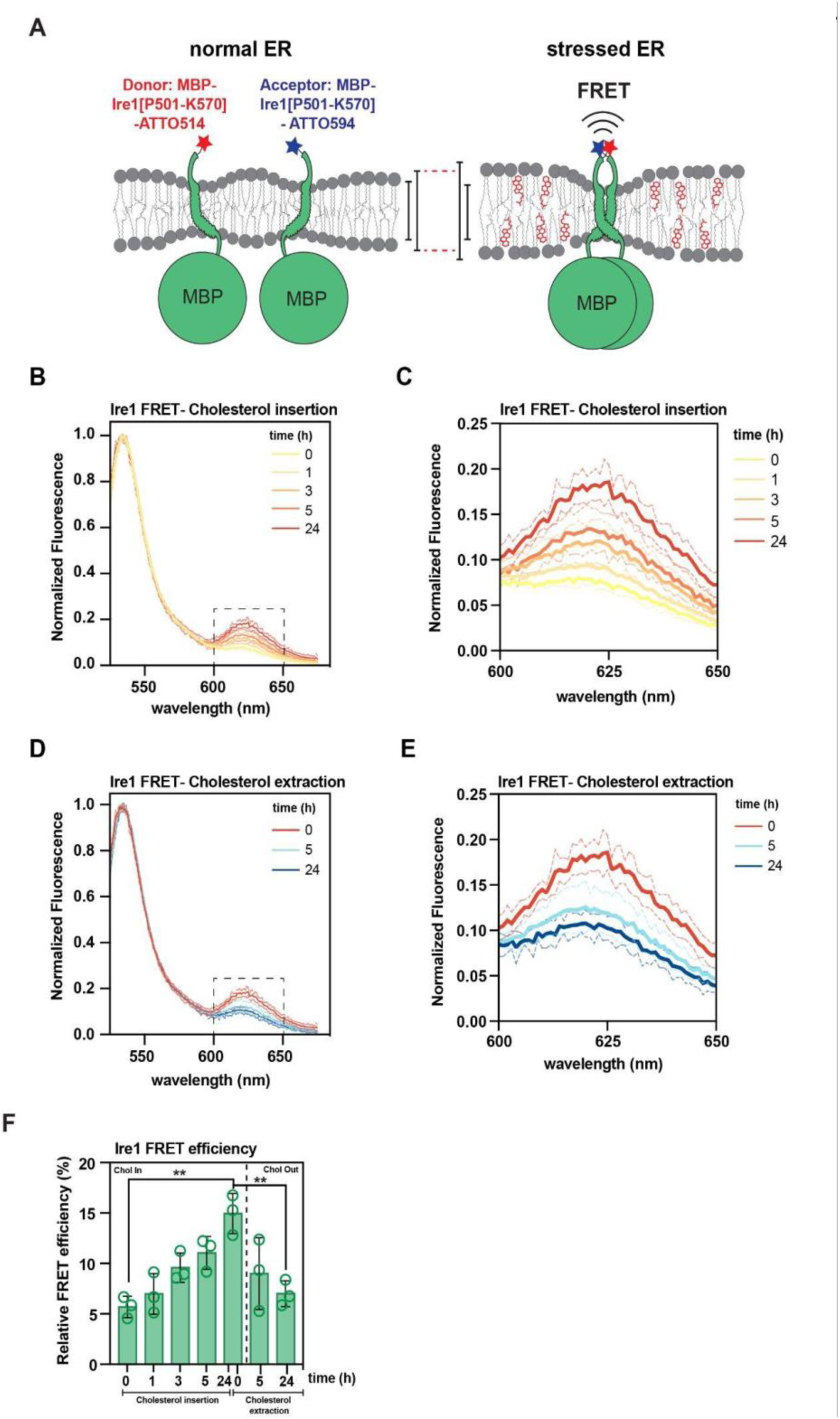
Cholesterol insertion into proteoliposomes induces Ire1 oligomerization in a reversible manner. **(A)** Ire1 oligomerization model in response to induced lipid bilayer stress *in vitro*. The reconstituted Ire1 transmembrane domain (Ire1^aa501-570^, C552S+C) is labeled with an N-terminal MBP-tag and a C-terminal ATTO dye. For the donor construct an ATTO514 is linked at the C terminal cysteine while in the acceptor construct an ATTO-594 is linked at the C terminal cysteine. Donor and/or acceptor constructs were reconstituted in cholesterol free 50% POPC 50% DOPC liposomes (normal ER). In turn, cholesterol was gradually inserted into the membrane inducing Ire1 oligomerization (stressed ER). Transverse membrane thickness of each model is represented by black bars. **(B)** Fluorescence emission spectra were recorded upon donor excitation (Ex: 514 nm; Em: 525–675 nm; slit width: 3 nm) and normalized to the maximal donor emission at indicated times during cholesterol delivery via cholesterol-loaded mβCD. Spectra are plotted as mean of three independent reconstitutions (n = 3; mean±SD). **(C)** Normalized fluorescence emission showing the FRET shoulder at several time points during 24 hours of cholesterol insertion using cholesterol-loaded mβCD. **(D)** Fluorescence emission spectra were recorded upon donor excitation (Ex: 514 nm; Em: 525–675 nm; slit width: 3 nm) and normalized to the maximal donor emission at indicated times during 24 hours of cholesterol extraction via empty mβCD. Spectra are plotted as mean of three independent reconstitutions (n = 3; mean±SD). **(E)** Normalized fluorescence emission showing the FRET shoulder at the indicated times during 24 hours of cholesterol extraction using empty mβCD. **(F)** The relative FRET efficiency was derived from the fluorescence spectra in (B) and (D) and plotted as mean of three independent reconstitutions (n = 3; mean±SD). A two-tailed, unpaired *t*-test was performed to test for statistical significance (\**p* < 0.05, \*\**p* < 0.01, \*\*\**p* < 0.001).

In contrast to classical reconstitution schemes, our mβCD-based transfer of cholesterol also allows for removing cholesterol from proteoliposomes. This reversibility also provides a means to distinguish between a functional, cholesterol-triggered oligomerization and an irreversible aggregation of a transmembrane protein from an unsuccessful reconstitution. After cholesterol was delivered to proteoliposomes containing Ire1-based donor and acceptor constructs (Fig. 4D, 0h), we removed it again by placing the dialysis cassette in a new bath containing empty mβCD. The removal of cholesterol from the proteoliposomes show reestablished membrane compressibility and dissociation of Ire1 dimers. Indeed, the relative FRET efficiency was substantially lower after 5 h or cholesterol removal and even more so after 24 h (Fig. 4D,E). Hence, using a dialysis-based sterol exchange setup combined with FRET we demonstrate the potential of this approach to manipulate the behavior of reconstituted transmembrane proteins in a controlled and reversible manner. Tuning sterol levels and membrane compressibility in preformed proteoliposomes could become a useful and widely used tool to study the impact of the lipid bilayer on membrane protein structure, dynamics, and function.

## Discussion

We have implemented an easy-to-use experimental setup to reversibly modify sterol levels in pre-existing (proteo)liposomes. Because dialysis does not require expensive instrumentation, this approach is broadly accessible to virtually every biophysical, pharmaceutical or biochemical laboratory. Sterol transfer to and from proteo(liposomes) is mediated by mβCD shuttling between two compartments separated by a dialysis membrane. Throughout sterol exchange, the (proteo)liposomes are retained in their compartment, thereby facilitating easy recovery, straightforward buffer exchange, and quantitative mβCD removal whenever necessary. Our proof-of-principle experiments with liposomes and proteoliposomes show that cholesterol delivery increases lipid packing as shown by C-Laurdan spectroscopy (Fig. 2, 3) and decreases membrane compressibility as suggested by the membrane-driven dimerization of a membrane property sensor module derived from Ire1 (Fig. 4) (4, 30–32).

The dialysis setup is versatile and provides several advantages: 1) Membrane material can be recovered with excellent yields. 2) Donor and acceptor liposomes remain separated throughout the experiment, thereby preventing undesired membrane fusion and facilitating a parallel, spectroscopic characterization of both samples (Fig. 3E-F). 3) Sterol-rich membrane environments can be established in pre-existing proteoliposomes, which often is challenging to achieve using ‘standard’ reconstitution procedures. 4) The gradual delivery of sterol is both time- and cost-effective: A single transmembrane protein reconstitution in a sterol-free membrane provides sufficient material for a whole set of samples covering a broad range of sterol concentrations and membrane compressibilities (Fig. 4). 5) The versatile dialysis setup is applicable not only to liposomes, but also to other nanodisc or complex biomembranes such as exosomes, thereby widening the scope of potential applications.

Nevertheless, the dialysis-based setup has limitations and not all possible applications have been explored. 1) Lipid delivery to liposomes (Fig. 2, 3) and proteoliposomes (Fig. 4) takes hours. Only sufficiently stable transmembrane proteins that ‘survive’ the time of dialysis can be suitably studied using this approach. 2) The specificity of mβCD does not guarantee the exclusive transfer of sterols, and some level of glycerophospholipid transfer cannot be ruled out. In fact, mβCD is known to bind and exchange glycerophospholipids at high concentrations (34, 45, 52–54), but less so at the concentrations used in this study. Yet, it will be important to quantify lipid recovery especially when handling (proteo)liposomes with complex lipid compositions. 3) While being theoretically useful for generating asymmetric liposomes, the cyclodextrin-mediated transfer of glycerophospholipids is too slow under the here established conditions (data not shown). A significant portion of glycerophospholipids delivered to the outer leaflet of an acceptor (proteo)liposome would passively flip to the luminal leaflet in the time of dialysis.

One of the key challenges in studying the structure, dynamics, and function of transmembrane proteins in complex, native-like membrane environments arises from the heterogeneous distribution of membrane components in (proteo)liposomes, which can be accentuated by the presence of sterols (15, 19–22). The mβCD-mediated delivery of sterols to pre-existing (proteo)liposomes, which is best accomplished in a dialysis setup, provides a potential work-around scheme to potentially overcome at least some of these issues. Because sterol delivery is reversible, we can distinguish between irreversible aggregation and reversible, membrane-based oligomerization as demonstrated for the sensory module of the membrane property sensor Ire1 (Fig. 4). We are convinced that this setup for lipid exchange can be applied to both model membranes and biomembranes by providing an on-demand delivery of sterols and/or phospholipids. After manipulation of the lipid composition, individual (proteo)liposome fractions can be removed from the dialysis setup and subjected to virtually any type of biophysical analysis including fluorescence spectroscopy, CD spectroscopy, native mass spectrometry, dynamic light scattering, electron-paramagnetic resonance spectroscopy, and cryo-electron microscopy. Hence, the dialysis-based setup for a mβCD-mediated lipid transfer will be a valuable tool to characterize the structure and function of membrane proteins in different lipid environments.

## Materials and methods

### Production of multilamellar vesicles

Liposomes of defined compositions were generated by mixing 1,2-dioleoyl-*sn*-glycero-3-phosphocholine (DOPC),1-palmitoyl-2-oleoyl-*sn*-glycero-3-phosphocholine (POPC) and cholesterol (25 mg/ml stock) or dehydroergosterol (DHE) (1 mg/ml stock) in chloroform. Those were: 1) 100% POPC; 2) 50% DOPC, 50% POPC; 3) X% cholesterol (100-X) % POPC; 4) X% cholesterol (50-X/2)% POPC (50-X/2)% DOPC; 5) X% DHE (100-X)% POPC. Typically, we prepared liposomes in batches containing 10 mM lipids. The solvent was evaporated under a stream of nitrogen in a heating block at temperatures well above the melting temperature of the lipids. For a complete removal of the solvent, the lipid film was placed in a desiccator under high vacuum for at least 1h at RT. For forming multilamellar vesicles (MVLs), the lipid cake was rehydrated with pre-warmed liposome buffer (20 mM HEPES pH 7.4, 150 mM NaCl, 5% glycerol) to reach the desired lipid concentration. Samples were agitated in a thermal mixer (60°C; 1200 rpm; 30 min) and then sonicated in a water bath for 20 min at 60 °C and at power setting 9 (VWR ultrasonic cleaner). The suspension of MLVs was used for protein reconstitution experiments at RT or snap-frozen with liquid nitrogen and stored at −80°C.

### Preparation of Large Unilamellar Vesicles (LUVs)

MLVs were subjected to 7 cycles of freeze-thawing in alternating liquid nitrogen and water bath at 40 °C prior to passing the sample with an extruder 31-times through membrane with 200 nm pore size. The glycerophospholipid concentration was determined by determining the amount of inorganic phosphate after lipid hydrolysis, following classic protocols (55).

### Liposome leakage assay

5(6)-Carboxyfluorescein (CF) loaded MLVs were prepared by rehydrating a lipid film (100% POPC) using the insight buffer (20 mM HEPES, 75 mM CF, pH 7.4). In turn, CF-loaded liposomes were subjected to 7 cycles of freeze thawing and extruded 10 times using a 200 µm membrane to produce LUVs. The suspension with CF-loaded LUVs was loaded onto a PD-10 column to remove free CF. For measuring the fluorescence of CF-loaded liposomes, a suspension at a lipid concentration of 20µM was placed in a 96-well plate, and the fluorescence was measured using a TECAN SPARK 20M plate reader (Ex: 492±5 nm; Em: 517±5 nm) over time. After the treatment of the liposomes with loaded mβCD or empty mβCD the CF fluorescence was observed over time. The addition of Triton X-100 to a final concentration of 0.1% (w/v) releases all CF by solubilizing the liposomes.

### Preparation of cholesterol-loaded mβCD

Cholesterol-loaded mβCD was prepared as described previously (34). Briefly, 132 mg of mβCD (100 µmol) and 11.9 mg of cholesterol (30.8 µmol) were dissolved in 600 µl methanol by rigorous mixing at room temperature. The solution was dried under a low of nitrogen. For full removal of the solvent from, loaded mβCD was transferred in a desiccator and vacuum was applied for 1 h. The material was resuspended with 2 ml phosphate-buffered saline (C_final_ = 50mM) and subjected to sonification in a water bath (37°C, full power, 3 min) and then incubated in a shaker at 37°C overnight.

### Cholesterol exchange setup

1 ml of cholesterol-loaded mβCD is diluted 20-fold in liposome buffer (20 mM HEPES pH 7.4, 150 mM NaCl, 5% glycerol) to yield the final volume of 21 ml for the outer bath. For cholesterol removal, 1 ml of a solution with empty mβCD (200 µmol) is used equivalently. Prior to their use, Spectra-Por® Float-A-Lyzer® G2 cassettes (100 kDa) are hydrated in liposome buffer, before they are placed in the outer bath (V = 21 ml) containing liposome buffer with either cholesterol-loaded or empty mβCD. Acceptor (proteo)liposomes are pipetted in the dialysis cassette (LUVs with 0.2 mM lipids; proteoliposomes with 0.6 mM lipids assuming 100% recovery during membrane protein reconstitution) after a small sample was taken as t = 0 min control. The outer bath is stirred with a magnetic stirrer at 270 rpm (VELP Scientifica-F203A0178). The dialysis setup was protected from daylight, whenever fluorescent molecules were used.

### C-Laurdan spectroscopy

To measure lipid packing, we used the solvatochromic dye 6-dodecanoyl-2-[N-methyl-N-(carboxymethyl)amino]naphthalene (C-Laurdan) (29) (Bio-Techne GmbH,7273). Fluorescence spectra were recorded (Ex: 375 ± 5 nm; Em: 400 – 530 nm; Em slit width: 5 nm) in a TECAN SPARK 20M plate reader at 30°C. Prior to the addition of C-laurdan, a first spectrum was recorded as scattering control. C-Lauran was added to the suspension at C-Laurdan-to-lipid ratio 1:500. After 3 min of equilibration, the C-Laurdan fluorescence spectrum was recorded (Ex: 375 ± 5 nm; Em: 400 – 530 nm; Em slit width: 5 nm). The C-Laurdan fluorescence emission spectrum was corrected, by subtracting the recorded intensities of the scattering control. The GP was calculated from the corrected C-Laurdan fluorescence emission spectrum by first integrating the intensities between 400 and 460 nm (*I*_Ch1_), and 470 and 530 nm (*I*_Ch2_), and then following the following equation:

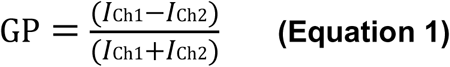

### Dynamic light scattering

Liposomes and other samples were subjected in re-usable Quartz microcuvettes (3×3 mm; 8,5 mm - 105. 251-QS, Hellma Analytics, Germany) to a Zetasizer Nano S (Malvern Panalytical, Worcestershire, UK) and analyzed for their dynamic light scattering (DLS) after 60s of equilibration at 25°C with the settings RI = 1,45 and at Abs = 0.001. All samples were measured three times (11 runs of 10s per measurement), with the attenuator position automatically optimized for each measurement. Data analysis was performed using the Zetasizer software version 7.13.

### DHE exchange experiments

1.5 ml of a solution containing 200 µmol empty mβCD was diluted 14-fold in liposome buffer (20 mM HEPES pH 7.4, 150 mM NaCl, 5% glycerol) to yield the final volume of 21 ml for the outer bath. Prior to their use, Spectra-Por® Float-A-Lyzer® G2 cassettes (100 kDa) are hydrated in liposome buffer. 1 ml of a suspension POPC-based liposomes containing 2 mol% DHE (200 µmol total lipid) were pipetted into the dialysis cassette with a 100 kDa molecular weight cutoff. Non-fluorescent, POPC-based acceptor liposomes (400 µmol total lipid) were added to the outer bath of the dialysis setup. At t = 0 min, just before the dialysis cassette was placed in the outer bath 5.7 µl was retrieved as a sample from inside the cassette and 120 µl was taken from the outer bath. After adjusting both samples to 120 µl using liposome buffer (20 mM HEPES pH 7.4, 150 mM NaCl, 5 % (w/v) glycerol) they were subjected to a quartz cuvette (10×2 mm; 15 mm-105.250-QS, Hellma Analytics, Germany). DHE was quantified using its fluorescence emission (Ex: 324±4 nm; Em: 394±4 nm) in a FluoroMax® 4 (Horiba, Japan) at 25°C. All fluorescence data were corrected for scattering using control spectra with identical lipid concentration, but lacking DHE.

### Bacterial cultivation for protein purification

*Escherichia coli* BL21 pLysS carrying the expression vector (pRE982) encoding for a fusion of the maltose-binding protein (MBP) and the transmembrane region of Ire1 (P501-K570) from *Saccharomyces cerevisiae* were cultivated in 50 ml of LB medium (containing ampicillin and chloramphenicol) at 37°C for 16 to 18 hours under continuous shaking. The overnight culture was used to inoculate 2 l of LB medium supplemented with 0.2% glucose, ampicillin, and chloramphenicol to an OD_600_ of 0.05 (37°C). The expression was induced with 0.3 mM IPTG at an OD_600_ of 0.6. After 3h at of cultivation at 37°C, the cells were harvested by centrifugation (3,000 x g, 20 min, 4°C), and the cell pellet was washed with ice-cold column buffer (50 mM HEPES pH 7.0, 150 mM NaCl, 1 mM EDTA, pH 8.0) and cells were harvested by centrifugation (3,000 x g, 20 minutes, 4°C). Pellets were stored at −20°C.

### Extraction, purification and labeling of a membrane property sensor

Typically, a cell pellet from a two liter bacterial culture was thawed on ice. Cells were resuspended in 45 ml ice-cold lysis buffer (50 mM HEPES pH 7.0, 150 mM NaCl, 1 mM EDTA pH 8.0, 50 mM 1-O-n-Octyl-β-D-glucopyranoside (OG)) containing 10 mg/ml chymostatin, 10 mg/ml antipain, 10 mg/ml pepstatin, 10 mM TCEP, 25 units/ml Benzonase® (Merck). The suspension was sonified with a VS 70T probe and SONOPULS HD 2070 (Bandelin) using 30% power and 70% duty in six cycles of 30s sonication each followed by an intermission of 30 s. The resulting lysates were rotated for 30 min at 4°C. Cell debris was removed by ultracentrifugation (100,000xg, 30 min, 4°C) using a Type 70 Ti rotor (Beckmann Coulter). The supernatant was transferred onto pre-equilibrated amylose beads in column buffer (50 mM HEPES pH 7.0, 150 mM NaCl, 1 mM EDTA-NaOH pH 8.0) and incubated under constant agitation for 60 min. The suspension of amylose beads was then distributed to two gravity columns. Each column was washed twice with 20 ml of degassed lysis buffer (50 mM HEPES pH 7.0, 150 mM NaCl, 1 mM EDTA pH 8.0, 50 mM OG). Next, 1.2 ml of labeling solution containing either ATTO514, ATTO594, or NEM at a concentration of 0.25 mM in lysis buffer (50 mM HEPES pH 7.0, 150 mM NaCl, 1 mM EDTA pH 8.0, 50 mM OG) was added and incubated with the beads and the bound protein overnight at 4°C under constant agitation. The next day, each column was washed three times with 20 ml lysis buffer. The labeled protein was eluted with 3 x 2 ml elution buffer (50 mM HEPES pH 7.0, 150 mM NaCl, 1 mM EDTA pH 8.0, 50 mM OG, 10 mM maltose, 10% (w/v) glycerol) each after a 5 min incubation on the column. The pooled eluate was then concentrated to a final volume of 600 µl using a spin concentrator with a 30 kDa molecular weight cutoff, snap-frozen in liquid nitrogen, and stored at −80°C for later use. For further purification, the protein was subjected to size exclusion chromatography. A protein aliquot was thawed at 23±1 °C and centrifuged (20,000 x g, 10 min, 4°C) to remove potential protein aggregates. 500 µl from the supernatant were loaded with 0.5 ml min^−1^ onto a Superdex® Increase 200 column equilibrated with gel filtration buffer (20 mM HEPES pH 7.4, 150 mM NaCl, 50 mM OG) and 250 µl fractions were collected. Protein-containing fractions were pooled and adjusted to a final glycerol concentration of 10% (w/v).

### Reconstitution of a membrane property sensor in liposomes

For each reconstitution, 200 µl of MLVs (10 mM stock) mixed with 20 mM HEPES pH 7.4, 150 mM NaCl, and 37.5 mM OG and agitated on a rotor for 10 min for complete lipid solubilization. Glycerol and SDS were added to reach a final concentration of 7% (w/v) and 0.3 mM in the reconstitution mix, respectively. Fluorescent (and non-fluorescent) proteins were included at a protein-to-lipid ratio of 1:16000. The total volume of the reconstitution mix was 1 ml. After 10 min agitation, the mix was transferred in a dialysis cassette with a 10 kDa molecular weight cutoff and dialyzed against 1 l dialysis buffer (20 mM HEPES pH 7.4, 150 mM NaCl, 5% glycerol). 400 mg of methanol-activated, washed, and equilibrated SM-2 Bio-Beads were added to the outer bath of the dialysis setup to provide a sink for detergent molecules. After 1 h, the dialysis cassette was placed in a new bath with 1 l dialysis buffer (without SM-2 Bio-Beads) and dialyzed for 1 h. This step was repeated twice. As the final step, the cassette was dialyzed over night against dialysis buffer with 800 mg methanol-activated, washed, and equilibrated SM-2 Bio-Beads to remove the last traces of OG and to yield proteoliposomes with the Ire1-based membrane property sensor.

### Determining the relative Förster Resonance Energy Transfer (FRET) efficiency

Ire1^NEM^, Ire1^ATTO514^ and Ire1^ATTO594^ constructs were used as unlabeled control, fluorescence donor and acceptor, respectively. Fluorescence emission spectra were recorded in proteoliposome buffer (20 mM HEPES pH 7.4, 150 mM NaCl, 7% (w/v) glycerol) and detergent solution (20 mM HEPES pH 7.4, 150 mM NaCl, 7% (w/v) glycerol, 50 mM OG, 4 mM SDS). Fluorescence spectra (Ex: 514±3 nm; Em: 512-800 nm; Em slid width: 3 nm) were recorded with an integration time to 0.1 s (FluoroMax® 4, Horiba, Japan) from 120 µl samples (0.6 mM lipid) in a 10×2 mm quartz cuvette (105.250-QS, Hellma Analytics, Germany) at 30°C. As the bleed-through for both the donor and acceptor fluorescence was low, and because only semi-quantiative information was required, we determined a ratiometric FRET (relative FRET: *E*_rel_). The fluorescence spectra were normalized to the highest emission (around 535 nm). Normalized spectra from a sample containing only the donor Ire1^ATTO514^ construct were subtracted from normalized spectra containing both Ire1^ATTO514^ and Ire1^ATTO594^ (donor and acceptor). The relative FRET efficiency was calculated as follows

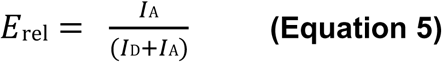

with I_A_ = maximum acceptor emission intensity at ≈ 620 nm, and I_D_ = maximum donor emission intensity at ≈ 535 nm.

## Author contributions

C.A. designed experiments, provided supervision performed research, analyzed data, and wrote the first draft of the manuscript. E.R. performed research and analyzed data. J.H. performed research, analyzed data, and provided technical and administrative support. M.F.R. contributed analytical tools and performed research. R.E. designed research, provided supervision, analyzed data, and wrote the manuscript.

## Acknowledgements

The authors wish to thank Alexander von der Malsburg for critical reading of the manuscript and John Reinhard for fruitful discussions in the early phase of the project. We a particularly grateful to Toni Radanović for his meticulous efforts in establishing new reconstitution protocols for Ire1. This work was funded by the Deutsche Forschungsgemeinschaft in the framework of the SFB1027 to R.E., and by the European Research Council under the European Union’s Horizon 2020 research and innovation program (grant agreement no. 866011) to R.E..

## Supplementary figures

**Figure S1:**
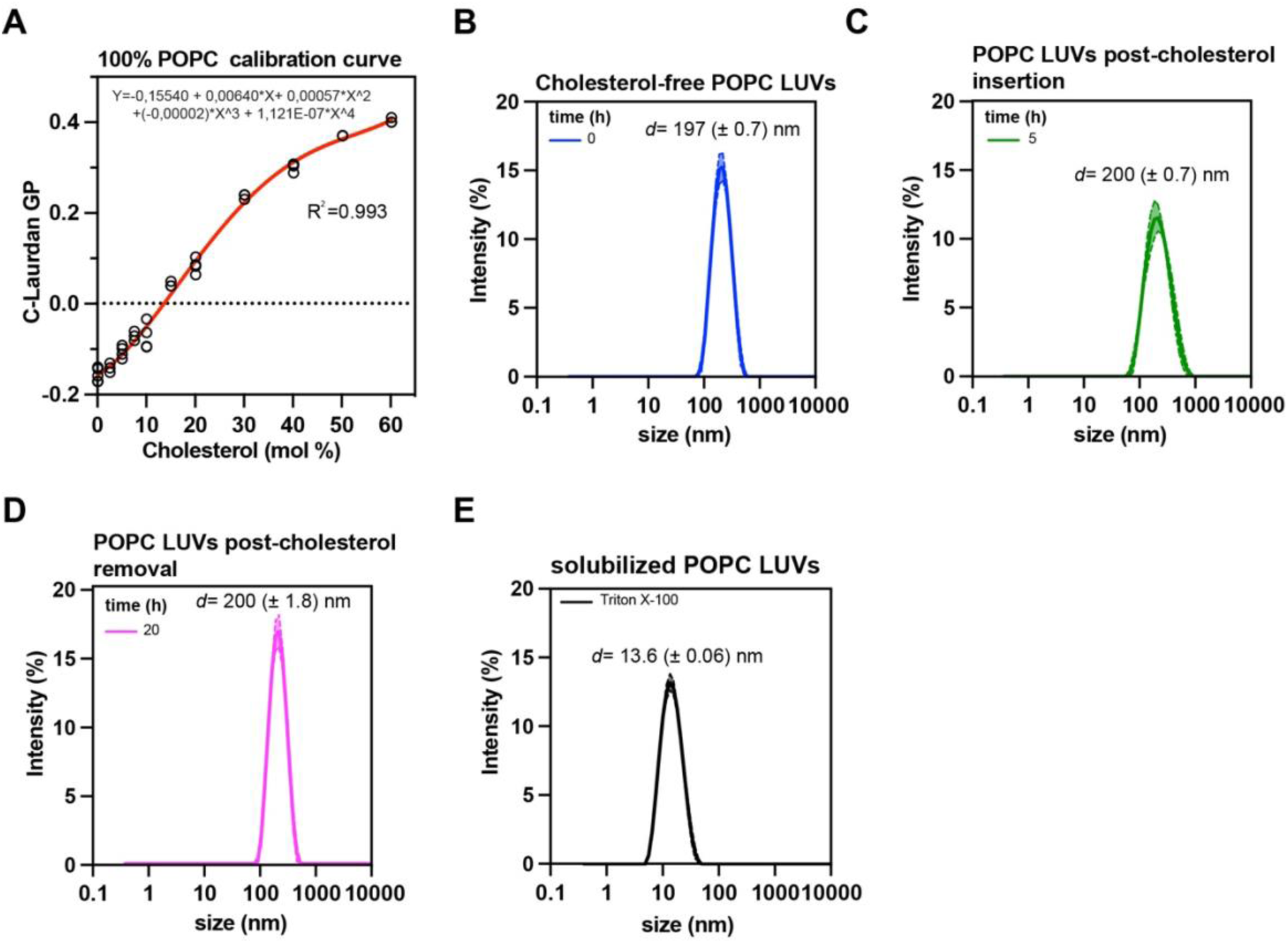
**(A)** The C-Laurdan GP of POPC-based liposomes containing different concentrations of cholesterol was determined and used to generate a calibration curve. A polynomial function was used to fit the experimental data. Each data point represents one of three independent experiments (n = 3). **(B-E)** The intensity-weighted particle size distribution of POPC-based liposomes was determined by DLS. Three technical replicates from a single LUV preparation are shown. The average particle size (or Z-average) is indicated as (*d*±SD). The individual samples are (B) liposomes prior to cholesterol delivery, (C) after 5 h of cholesterol delivery, (D) after subsequent cholesterol removal, and (E) after complete solubilization through the addition of with Triton X-100 (E).

**Figure S2:**
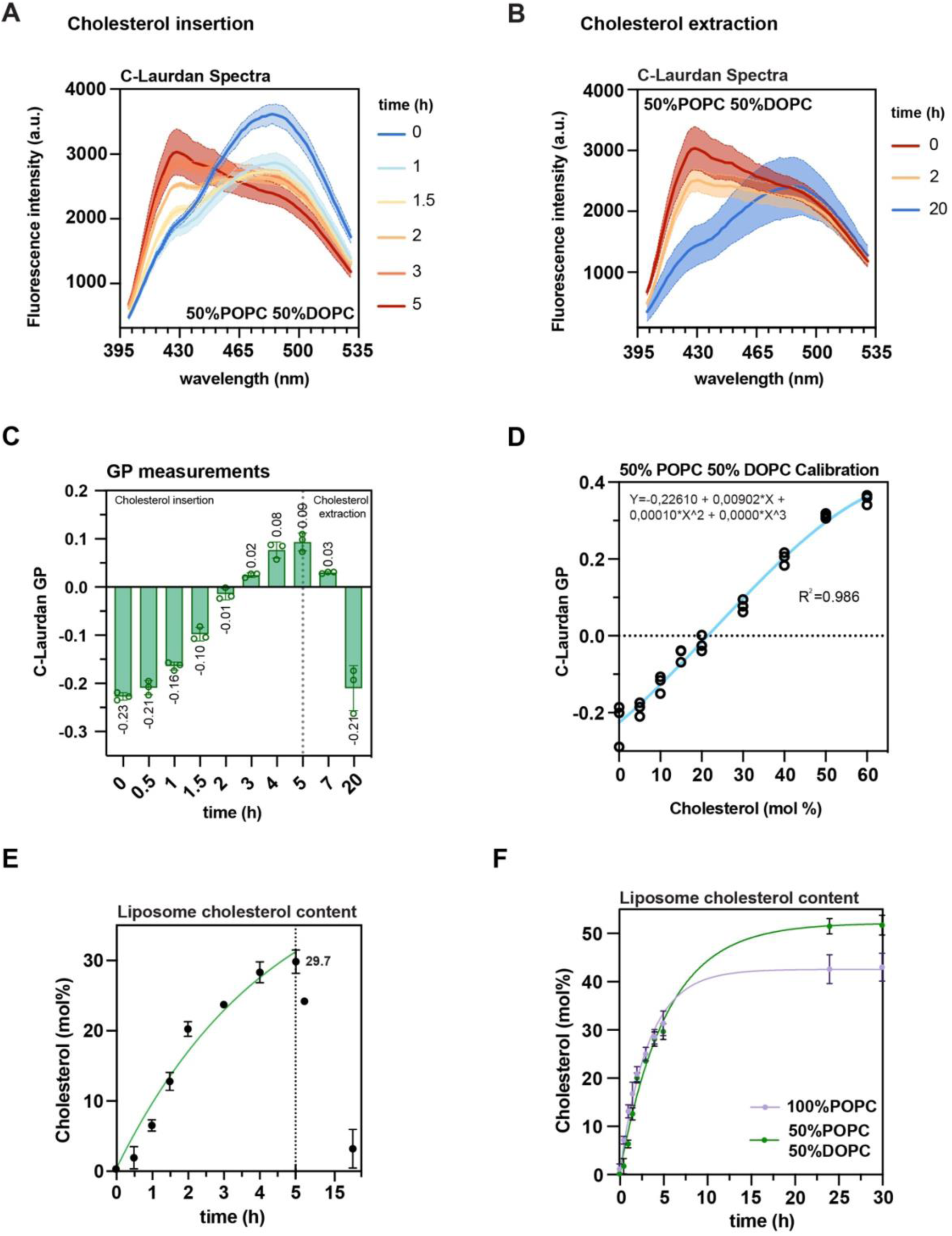
Cholesterol delivery and removal to and from LUVs composed of equimolar mix of POPC and POPC. **(A)** C-Laurdan fluorescence emission spectra (Ex: 375±5 nm; Em slit width: 5 nm) of POPC:DOPC-based liposomes retrieved from the dialysis cassette after indicated times of cholesterol delivery using mβCD. Data are from three independent experiments (n = 3;mean±SD). **(B)** C-Laurdan fluorescence emission spectra upon cholesterol removal using empty mβCD in the outer bath of the dialysis setup. Data are from three independent experiments (n = 3;mean±SD). **(C)** C-Laurdan GP of 50% POPC 50% DOPC LUVs during the exchange (20h) at room temperature. The dotted vertical line indicates the shift from cholesterol insertion to cholesterol extraction at t = 5h. Shown are independent triplicates (n = 3; mean±SD). **(D)** The C-Laurdan GP of POPC:DOPC-based liposomes containing different concentrations of cholesterol was determined and used to generate a calibration curve. A polynomial function was used to fit the experimental data. Each data point represents one of three independent experiments (n = 3). **(E)** Cholesterol concentration (in mol%) derived from a standard curve during cholesterol delivery and removal with POPC:DOPC-based liposomes. The dotted vertical line (at t = 5h) indicates the exchange of the outer batch and a switch from cholesterol insertion to cholesterol extraction. The purple solid line fitted to the data was obtained using a one phase association model (Prism 10). Shown are data from three independent experiments (n = 3; mean±SD). **(F)** Comparison of cholesterol delivery rates into POPC-based (purple) and POPC:DOPC liposomes (green). Data in this graph are identical to the date from Fig. 2E and Fig. S2E with additional included timepoints (24h and 30h) used to determine the plateaus of the cholesterol exchange. The experimental data are derived from three independent experiments (n = 3; mean±SD) and fitted using one phase association model (Prism 10).

**Figure S3:**
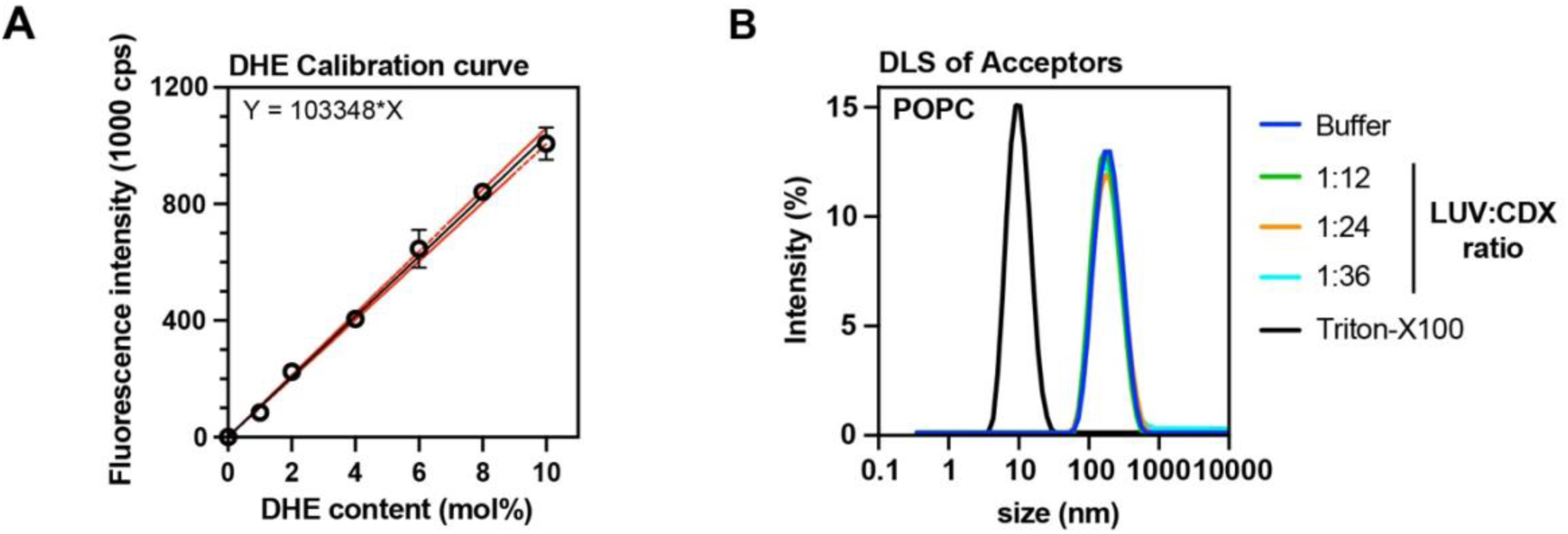
DHE calibration series for DHE content calculation. **(A)** Representative changes of DHE fluorescence intensities (ex: 324 nm; em: 394 nm; slit sizes: 4nm) in liposomes starting from 0% DHE content up to 10% DHE (9.5 µM lipid). The solid black line represents a linear regression. The data are derived from three independent experiments (n = 3; mean±SD). **(B)** The intensity-weighted particle size distribution of POPC-based liposomes was determined by DLS. The experimental conditions and the lipid:cyclodextrin ratio have no apparent impact on the size of LUVs. In the presence of Triton X-100, liposomes are solubilized.

**Figure S4:**
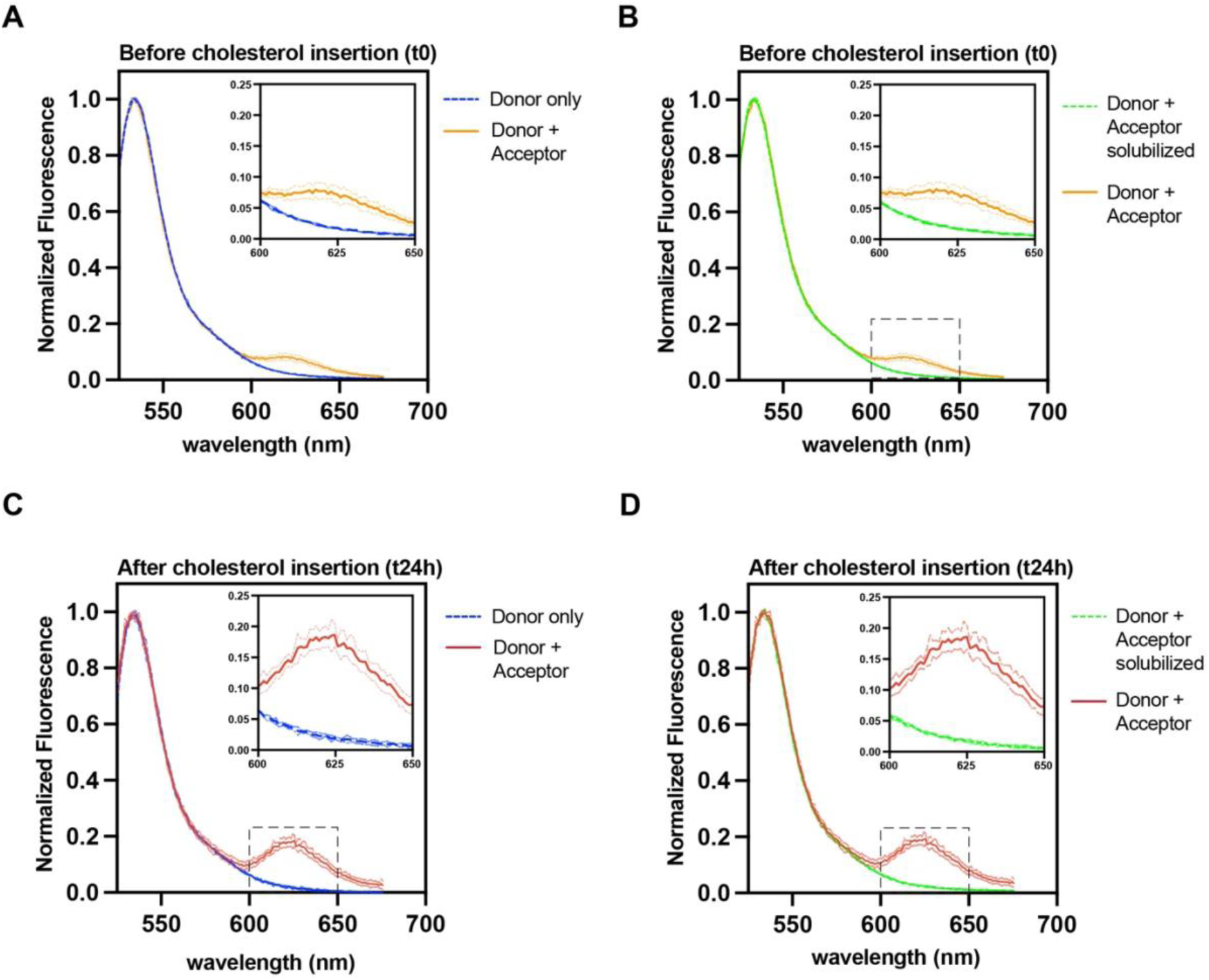
Validating the impact of cholesterol on the oligomerization of Ire1. **(A)** Fluorescence emission spectra were recorded upon donor excitation (Ex: 514 nm; Em: 525–675 nm; slit width: 3 nm) Fluorescence emission spectra (em: 525–675 nm) recorded upon excitation of the ATTO514 labeled donor (ex: 514 nm) and normalized to the maximal donor emission at t0 before cholesterol insertion in the absence (Donor only) and presence of acceptor (Donor+Acceptor). (**B)** Fluorescence emission spectra (em: 525–675 nm) were recorded upon excitation of the ATTO514 labeled donor (ex: 514 nm) and normalized to the maximal donor emission at t0 before cholesterol insertion (Donor+Acceptor) and after sample solubilization with detergent (Donor+Acceptor solubilized). **(C)** Fluorescence emission spectra (em: 525–675 nm) recorded upon excitation of the ATTO514 labeled donor (ex: 514 nm) and normalized to the maximal donor emission after cholesterol insertion (t24h) in the absence (Donor only) and presence of acceptor (Donor+Acceptor). **(D)** Fluorescence emission spectra (em: 525–675 nm) were recorded upon excitation of the ATTO514 labeled donor (ex: 514 nm) and normalized to the maximal donor emission after cholesterol insertion (t24h) (Donor+Acceptor) and s sample solubilization with detergent (Donor+Acceptor solubilized). All spectra (A to D) are plotted as mean of three independent reconstitutions (n = 3; mean±SD).

## References

1. Andersen, O.S., and R.E. Koeppe 2nd. 2007. Bilayer thickness and membrane protein function: an energetic perspective. Annu. Rev. Biophys. Biomol. Struct. 36:107–130.

2. Dowhan, W., H. Vitrac, and M. Bogdanov. 2019. Lipid-assisted membrane protein folding and topogenesis. Protein J. 38:274–288.

3. Levental, I., and E. Lyman. 2023. Regulation of membrane protein structure and function by their lipid nano-environment. Nat. Rev. Mol. Cell Biol. 24:107–122.

4. Renne, M.F., and R. Ernst. 2023. Membrane homeostasis beyond fluidity: control of membrane compressibility. Trends Biochem. Sci. 48:963–977.

5. Maxfield, F.R., and I. Tabas. 2005. Role of cholesterol and lipid organization in disease. Nature. 438:612–621.

6. Heberle, F.A., and G.W. Feigenson. 2011. Phase separation in lipid membranes. Cold Spring Harb. Perspect. Biol. 3:a004630–a004630.

7. Sezgin, E., I. Levental, S. Mayor, and C. Eggeling. 2017. The mystery of membrane organization: composition, regulation and roles of lipid rafts. Nat. Rev. Mol. Cell Biol. 18:361–374.

8. Frallicciardi, J., J. Melcr, P. Siginou, S.J. Marrink, and B. Poolman. 2022. Membrane thickness, lipid phase and sterol type are determining factors in the permeability of membranes to small solutes. Nat. Commun. 13:1605.

9. Doole, F.T., T. Kumarage, R. Ashkar, and M.F. Brown. 2022. Cholesterol stiffening of lipid membranes. J. Membr. Biol. 255:385–405.

10. Doktorova, M., J.L. Symons, X. Zhang, H.-Y. Wang, J. Schlegel, J.H. Lorent, F.A. Heberle, E. Sezgin, E. Lyman, K.R. Levental, and I. Levental. 2023. Cell membranes sustain phospholipid imbalance via cholesterol asymmetry. bioRxiv. 2023.07.30.551157.

11. Rawicz, W., B.A. Smith, T.J. McIntosh, S.A. Simon, and E. Evans. 2008. Elasticity, strength, and water permeability of bilayers that contain raft microdomain-forming lipids. Biophys. J. 94:4725–4736.

12. Killian, J.A. 1998. Hydrophobic mismatch between proteins and lipids in membranes. Biochim. Biophys. Acta. 1376:401–415.

13. Kaiser, H.-J., A. Orłowski, T. Róg, T.K.M. Nyholm, W. Chai, T. Feizi, D. Lingwood, I. Vattulainen, and K. Simons. 2011. Lateral sorting in model membranes by cholesterol-mediated hydrophobic matching. Proc. Natl. Acad. Sci. U. S. A. 108:16628–16633.

14. Milovanovic, D., A. Honigmann, S. Koike, F. Göttfert, G. Pähler, M. Junius, S. Müllar, U. Diederichsen, A. Janshoff, H. Grubmüller, H.J. Risselada, C. Eggeling, S.W. Hell, G. van den Bogaart, and R. Jahn. 2015. Hydrophobic mismatch sorts SNARE proteins into distinct membrane domains. Nat. Commun. 6:5984.

15. Menon, I., T. Sych, Y. Son, T. Morizumi, J. Lee, O.P. Ernst, G. Khelashvili, E. Sezgin, J. Levitz, and A.K. Menon. 2024. A cholesterol switch controls phospholipid scrambling by G protein-coupled receptors. J. Biol. Chem. 300:105649.

16. Li, D., C. Rocha-Roa, M.A. Schilling, K.M. Reinisch, and S. Vanni. 2024. Lipid scrambling is a general feature of protein insertases. Proc. Natl. Acad. Sci. U. S. A. 121:e2319476121.

17. Nagle, J.F. 2013. Introductory lecture: basic quantities in model biomembranes. Faraday Discuss. 161:11–29; discussion 113-50.

18. Harayama, T., and H. Riezman. 2018. Understanding the diversity of membrane lipid composition. Nat. Rev. Mol. Cell Biol. 19:281–296.

19. Larsen, J., N.S. Hatzakis, and D. Stamou. 2011. Observation of inhomogeneity in the lipid composition of individual nanoscale liposomes. J. Am. Chem. Soc. 133:10685–10687.

20. Elizondo, E., J. Larsen, N.S. Hatzakis, I. Cabrera, T. Bjørnholm, J. Veciana, D. Stamou, and N. Ventosa. 2012. Influence of the preparation route on the supramolecular organization of lipids in a vesicular system. J. Am. Chem. Soc. 134:1918–1921.

21. Cliff, L., R. Chadda, and J.L. Robertson. 2020. Occupancy distributions of membrane proteins in heterogeneous liposome populations. Biochim. Biophys. Acta Biomembr. 1862:183033.

22. Sych, T., J. Schlegel, H.M.G. Barriga, M. Ojansivu, L. Hanke, F. Weber, R. Beklem Bostancioglu, K. Ezzat, H. Stangl, B. Plochberger, J. Laurencikiene, S. El Andaloussi, D. Fürth, M.M. Stevens, and E. Sezgin. 2024. High-throughput measurement of the content and properties of nano-sized bioparticles with single-particle profiler. Nat. Biotechnol. 42:587–590.

23. Ohtani, Y., T. Irie, K. Uekama, K. Fukunaga, and J. Pitha. 1989. Differential effects of alpha-, beta- and gamma-cyclodextrins on human erythrocytes. Eur. J. Biochem. 186:17–22.

24. Huang, Z., and E. London. 2013. Effect of cyclodextrin and membrane lipid structure upon cyclodextrin-lipid interaction. Langmuir. 29:14631–14638.

25. Li, G., J. Kim, Z. Huang, J.R. St Clair, D.A. Brown, and E. London. 2016. Efficient replacement of plasma membrane outer leaflet phospholipids and sphingolipids in cells with exogenous lipids. Proc. Natl. Acad. Sci. U. S. A. 113:14025–14030.

26. Li, G., S. Kakuda, P. Suresh, D. Canals, S. Salamone, and E. London. 2019. Replacing plasma membrane outer leaflet lipids with exogenous lipid without damaging membrane integrity. PLoS One. 14:e0223572.

27. Zidovetzki, R., and I. Levitan. 2007. Use of cyclodextrins to manipulate plasma membrane cholesterol content: evidence, misconceptions and control strategies. Biochim. Biophys. Acta. 1768:1311–1324.

28. Crini, G. 2014. Review: a history of cyclodextrins. Chem. Rev. 114:10940–10975.

29. Kim, H.M., H.-J. Choo, S.-Y. Jung, Y.-G. Ko, W.-H. Park, S.-J. Jeon, C.H. Kim, T. Joo, and B.R. Cho. 2007. A two-photon fluorescent probe for lipid raft imaging: C-laurdan. Chembiochem. 8:553–559.

30. Halbleib, K., K. Pesek, R. Covino, H.F. Hofbauer, D. Wunnicke, I. Hänelt, G. Hummer, and R. Ernst. 2017. Activation of the unfolded protein response by lipid bilayer stress. Mol. Cell. 67:673–684.e8.

31. Väth, K., C. Mattes, J. Reinhard, R. Covino, H. Stumpf, G. Hummer, and R. Ernst. 2021. Cysteine cross-linking in native membranes establishes the transmembrane architecture of Ire1. J. Cell Biol. 220.

32. Ernst, R., M.F. Renne, A. Jain, and A. von der Malsburg. 2024. Endoplasmic reticulum membrane homeostasis and the unfolded protein response. Cold Spring Harb. Perspect. Biol. 16:a041400.

33. Christian, A.E., M.P. Haynes, M.C. Phillips, and G.H. Rothblat. 1997. Use of cyclodextrins for manipulating cellular cholesterol content. J. Lipid Res. 38:2264–2272.

34. Cheng, H.-T., Megha, and E. London. 2009. Preparation and properties of asymmetric vesicles that mimic cell membranes: effect upon lipid raft formation and transmembrane helix orientation. J. Biol. Chem. 284:6079–6092.

35. Ruysschaert, T., A. Marque, J.-L. Duteyrat, S. Lesieur, M. Winterhalter, and D. Fournier. 2005. Liposome retention in size exclusion chromatography. BMC Biotechnol. 5:11.

36. Kaiser, H.-J., D. Lingwood, I. Levental, J.L. Sampaio, L. Kalvodova, L. Rajendran, and K. Simons. 2009. Order of lipid phases in model and plasma membranes. Proc. Natl. Acad. Sci. U. S. A. 106:16645–16650.

37. Steinkühler, J., E. Sezgin, I. Urbančič, C. Eggeling, and R. Dimova. 2019. Mechanical properties of plasma membrane vesicles correlate with lipid order, viscosity and cell density. Commun. Biol. 2:337.

38. Yeagle, P.L., and J.E. Young. 1986. Factors contributing to the distribution of cholesterol among phospholipid vesicles. J. Biol. Chem. 261:8175–8181.

39. Leventis, R., and J.R. Silvius. 2001. Use of cyclodextrins to monitor transbilayer movement and differential lipid affinities of cholesterol. Biophys. J. 81:2257–2267.

40. Niu, S.-L., and B.J. Litman. 2002. Determination of membrane cholesterol partition coefficient using a lipid vesicle-cyclodextrin binary system: effect of phospholipid acyl chain unsaturation and headgroup composition. Biophys. J. 83:3408–3415.

41. Tsamaloukas, A., H. Szadkowska, P.J. Slotte, and H. Heerklotz. 2005. Interactions of cholesterol with lipid membranes and cyclodextrin characterized by calorimetry. Biophys. J. 89:1109–1119.

42. Pan, J., S. Tristram-Nagle, and J.F. Nagle. 2009. Effect of cholesterol on structural and mechanical properties of membranes depends on lipid chain saturation. Phys. Rev. E Stat. Nonlin. Soft Matter Phys. 80:021931.

43. Lönnfors, M., J.P.F. Doux, J.A. Killian, T.K.M. Nyholm, and J.P. Slotte. 2011. Sterols have higher affinity for sphingomyelin than for phosphatidylcholine bilayers even at equal acyl-chain order. Biophys. J. 100:2633–2641.

44. Nyholm, T.K.M., S. Jaikishan, O. Engberg, V. Hautala, and J.P. Slotte. 2019. The affinity of sterols for different phospholipid classes and its impact on lateral segregation. Biophys. J. 116:296–307.

45. Anderson, T.G., A. Tan, P. Ganz, and J. Seelig. 2004. Calorimetric measurement of phospholipid interaction with methyl-beta-cyclodextrin. Biochemistry. 43:2251–2261.

46. Reagle, T., Y. Xie, Z. Li, W. Carnero, and T. Baumgart. 2024. Methyl-β-cyclodextrin asymmetrically extracts phospholipid from bilayers, granting tunable control over differential stress in lipid vesicles. Soft Matter. 20:4291–4307.

47. Nasr, G., H. Greige-Gerges, S. Fourmentin, A. Elaissari, and N. Khreich. 2023. Cyclodextrins permeabilize DPPC liposome membranes: a focus on cholesterol content, cyclodextrin type, and concentration. Beilstein J. Org. Chem. 19:1570–1579.

48. Chen, R.F., and J.R. Knutson. 1988. Mechanism of fluorescence concentration quenching of carboxyfluorescein in liposomes: energy transfer to nonfluorescent dimers. Anal. Biochem. 172:61–77.

49. Walter, P., and D. Ron. 2011. The unfolded protein response: from stress pathway to homeostatic regulation. Science. 334:1081–1086.

50. Reinhard, J., L. Starke, C. Klose, P. Haberkant, H. Hammarén, F. Stein, O. Klein, C. Berhorst, H. Stumpf, J.P. Sáenz, J. Hub, M. Schuldiner, and R. Ernst. 2024. MemPrep, a new technology for isolating organellar membranes provides fingerprints of lipid bilayer stress. EMBO J. 43:1653–1685.

51. Covino, R., S. Ballweg, C. Stordeur, J.B. Michaelis, K. Puth, F. Wernig, A. Bahrami, A.M. Ernst, G. Hummer, and R. Ernst. 2016. A eukaryotic sensor for membrane lipid saturation. Mol. Cell. 63:49–59.

52. Ottico, E., A. Prinetti, S. Prioni, C. Giannotta, L. Basso, V. Chigorno, and S. Sonnino. 2003. Dynamics of membrane lipid domains in neuronal cells differentiated in culture. J. Lipid Res. 44:2142–2151.

53. Doktorova, M., F.A. Heberle, B. Eicher, R.F. Standaert, J. Katsaras, E. London, G. Pabst, and D. Marquardt. 2018. Preparation of asymmetric phospholipid vesicles for use as cell membrane models. Nat. Protoc. 13:2086–2101.

54. Krompers, M., and H. Heerklotz. 2023. A guide to your desired lipid-asymmetric vesicles. Membranes (Basel). 13.

55. Rouser, G., S. Fkeischer, and A. Yamamoto. 1970. Two dimensional then layer chromatographic separation of polar lipids and determination of phospholipids by phosphorus analysis of spots. Lipids. 5:494–496.

